# Kif11 overexpression rescues cognition, long-term potentiation, and spine defects in mouse and cell models of Alzheimer’s disease

**DOI:** 10.1101/2021.09.09.459627

**Authors:** Esteban M. Lucero, Ronald K. Freund, Noah R. Johnson, Breanna Dooling, Emily Sullivan, Olga Prikhodko, Md. Mahiuddin Ahmed, Mark L. Dell’Acqua, Heidi J. Chial, Huntington Potter

**Affiliations:** Department of Neurology, University of Colorado Anschutz Medical Campus, Aurora, CO, USA; University of Colorado Alzheimer’s and Cognition Center, University of Colorado Anschutz Medical Campus, Aurora, CO, USA; Linda Crnic Institute for Down Syndrome, University of Colorado Anschutz Medical Campus, Aurora, CO, USA; Department of Pharmacology, University of Colorado Anschutz Medical Campus, Aurora, CO, USA

**Keywords:** Alzheimer’s disease (AD), 5xFAD, Kinesin-5, Eg5, Kif11, microtubules, Alzheimer’s mouse model, neuronal structures, motor protein, learning and memory, amyloid beta (Aβ)

## Abstract

Competitive inhibition of kinesin motor proteins by amyloid-beta (Aβ) may contribute to alterations in the neuronal microtubule cytoskeleton that can disrupt plasticity mechanisms required for learning and memory, such as long-term potentiation (LTP), thus contributing to synaptic dysfunction and cognitive impairments associated with Alzheimer’s disease (AD). Here, we tested the hypothesis that overexpression of the microtubule motor protein KIF11 (Kinesin-5/Eg5) will rescue Aβ-mediated synaptic dysfunction and cognitive impairments. We found that overexpression of *Kif11* prevented spatial learning and LTP deficits in the 5xFAD mouse model of AD and rescued Aβ-mediated decreases in postsynaptic dendritic spine density in neuronal cultures. Together, these data suggest that KIF11 function is important for preserving synaptic structures and functions that are critical for learning and memory and for protection against Aβ-mediated loss of cognition in AD.

**Highlights:**

- Deficits in cognition and long-term potentiation in the 5xFAD mouse model of Alzheimer’s disease are prevented by *Kif11* overexpression.
- Aβ-mediated dendritic spine loss is blocked by *Kif11* overexpression.

## Introduction

The Alzheimer’s disease (AD) brain exhibits the characteristic pathology of amyloid deposits, comprised primarily of the amyloid-β (Aβ) peptide, and intracellular neurofibrillary tangles, comprised primarily of hyperphosphorylated forms of the microtubule associated protein tau. These hallmark pathological features of AD, in tandem with many reports elucidating the toxic effects of the Aβ peptide, have led to the development of the amyloid cascade hypothesis (Hardy and Higgins, 1992). This hypothesis postulates that the neuronal accumulation of the Aβ peptide, a cleavage product of the amyloid precursor protein (APP), initiates a cascade of molecular events that lead to the development and progression of the neuronal damage and cognitive deficits associated with AD. The toxic effects of Aβ induce microtubule instability that can be directly linked to cellular dysfunctions affecting synaptic plasticity, such as impaired organelle transport, decreased dendritic spine density, and altered cell cycle regulation that leads to chromosome mis-segregation (Spires et al., 2005, Golovyashkina et al., 2015, Wu et al., 2010, Wei et al., 2010, Umeda et al., 2015, Geller and Potter, 1999, Potter et al., 2019). Further highlighting the role of dysregulated microtubule stability in AD, pathways involved in microtubule organization have recently been linked to the susceptibility of neurons to AD (Roussarie et al., 2020). Additional studies showed that treatment with microtubule-stabilizing drugs can ameliorate AD phenotypes, including cognitive deficits, in model systems (Fernandez-Valenzuela et al., 2020, Zhang et al., 2018, Penazzi et al., 2016). Taken together, these findings demonstrate that the microtubule network plays a major role in establishing and maintaining neuronal structures and functions that are important for learning and memory and are disrupted in neurodegenerative diseases, such as AD (Dent, 2017).

In our previous search for potential mechanisms by which Aβ causes neurotoxicity, we discovered that the microtubule motor proteins KIF11 (Kinesin-5/Eg5), KIF4A, and MCAK/KIF2C are inhibited by soluble Aβ oligomers in a dose-dependent manner (Borysov et al., 2011). The inhibition of this select set of motor proteins leads to both microtubule and mitotic defects (Borysov et al., 2011). Kinetic analyses of KIF11, KIF4A, and MCAK motor function revealed the competitive nature of the inhibition by Aβ (Borysov et al., 2011). Of the kinesin motor proteins inhibited by Aβ, we were particularly interested in KIF11 due to a previously-reported association between a single nucleotide polymorphism (SNP) within the KIF11 haploblock region and both AD and type II diabetes, which is a major risk factor for AD (Feuk et al., 2005), as well as the finding that KIF11 expression is expressed in post-mitotic neurons where it contributes to neuronal structure and function (Freund et al., 2016, Freixo et al., 2018, Kahn et al., 2015). We also found that inhibition of rodent Kif11 by the specific chemical inhibitor monastrol in cell culture results in AD-like phenotypes that mimic Aβ toxicity, and that co-treatment of *ex vivo* hippocampal slices with monastrol and Aβ show that both Aβ and Kif11 act in the same pathway that leads to deficits in long-term potentiation (LTP) (Ari et al., 2014, Freund et al., 2016). These results suggest that Aβ-mediated inhibition of KIF11 in humans may contribute at least in part to the cognitive dysfunction observed in AD.

KIF11, the only member of the kinesin-5 family found in vertebrates, is a plus end-directed microtubule motor protein in mammalian cells that is best known for its role in mitosis where it contributes to bipolar spindle formation and provides the force required to separate the centrosomes and attached sister chromatids during cell division (Bodrug et al., 2020, Ferenz et al., 2010, Mann and Wadsworth, 2019, Blangy et al., 1995, Kapitein et al., 2005). In addition to its role in cell division, KIF11 is required for protein synthesis and serves as a link between ribosomes and microtubules during interphase in mammalian cells (Bartoli et al., 2011). KIF11 has also been shown to bind to the plus ends of microtubules and to enhance polymerization *in vitro* by stabilizing tubulin at the growing end of the microtubules, providing evidence that KIF11 regulates microtubule dynamics (Chen and Hancock, 2015).

An early study identified mouse *Kif11* expression in post-mitotic neurons (Ferhat et al., 1998), and later studies showed that Kif11 regulates axonal growth and contributes to dendritic architecture and spine formation, where inhibition, knockdown, or overexpression of *Kif11* led to abnormal neurite outgrowth (Freixo et al., 2018, Myers and Baas, 2007, Kahn et al., 2015, Nadar et al., 2008). In addition to its role in regulating neuronal morphology, the functional impact of Kif11 on neuronal microtubule dynamics and on interactions between microtubules has been shown to play a key role in regulating neuronal migration and growth cone dynamics (Falnikar et al., 2011, Nadar et al., 2008). In sum, KIF11 carries out key roles in neurons that derive from its microtubule interactions and contribute to neuronal cell morphology, development, migration, and function. These data highlight the importance of regulating KIF11 expression and function during the development of neuronal structures important for learning and memory.

Our previous reports showing that Aβ competitively inhibits KIF11 and leads to the breakdown of the microtubule network and to chromosome mis-segregation and aneuploidy (Ari et al., 2014, Borysov et al., 2011), and that inhibition of KIF11 by Aβ leads to deficits in LTP in *ex vivo* electrophysiology studies (Freund et al., 2016) highlight the importance of KIF11 for key neuronal functions in *in vitro* and *ex vivo* model systems. However, the role of KIF11 in learning and memory during normal aging and AD *in vivo* has yet to be elucidated.

Based on the known functions of KIF11 and on the cytoskeletal defects caused by Aβ toxicity, we hypothesized that Aβ-mediated inhibition of KIF11 that disrupts learning and memory may be prevented by increasing the expression of *KIF11* to preserve its key cellular functions. Specifically, we predicted that increased expression of *KIF11* would override its inhibition by Aβ and prevent or reduce cognitive deficits attributed to Aβ toxicity *in vivo*. To test our hypothesis, we generated a new mouse model derived by mating the 5xFAD Alzheimer’s mouse model (Oakley et al., 2006) to a mouse that overexpresses mouse *Kif11* (Castillo et al., 2007) to yield *Kif11*-overexpressing 5xFAD mice (5xFAD-Kif11OE). We then carried out behavioral, pathological, and electrophysiological studies to determine the effect(s) of increasing *Kif11* expression on Aβ-associated AD phenotypes. We also employed a primary neuron model of AD that, through transient transfection, overexpresses mouse *Kif11* and a human FAD mutant form of *APP* (*APPswe*) to interrogate the mechanisms that underlie *KIF11* and Aβ mediated effects on neuronal structure and function. Our results indicate that enhanced expression of *Kif11* obviates multiple deleterious effects of the Aβ peptide observed in 5xFAD mice and in primary rat neurons, and that increased levels of KIF11 protect against the effects of Aβ toxicity.

## Results

### Generating the 5xFAD-Kif11OE mouse model

In order to investigate the role of KIF11 in AD, we overexpressed mouse *Kif11* in the 5xFAD mouse model of AD. The 5xFAD transgenic mouse expresses the human *APP* gene harboring the “Swedish” double mutation (K670N/M671L), the “Florida” mutation (I716V), and the “London” mutation (V717I), and the human *PSEN1* gene harboring the M146L and L286V Familial/Early onset AD (FAD/EOAD) mutations. Together, these mutations result in the overproduction of Aβ and the deposition of neuronal Aβ plaques prior to six months of age (Bilkei-Gorzo, 2014, Hall and Roberson, 2012, Oakley et al., 2006),

The Kif11OE mouse was originally developed to determine whether overexpression of mouse *Kif11* leads to genomic instability and cancer (Castillo et al., 2007). In this model, *Kif11* overexpression led to mitotic spindle defects, increased genomic instability, higher rates of aneuploidy, and increased tumor development. In addition to these cancer-related phenotypes, Kif11OE mice also exhibit neurological abnormalities, megacystis, and dermatitis. However, the specific neurological phenotypes and effects of *Kif11* overexpression on learning and memory, have not been examined. By crossing the 5xFAD and Kif11OE strains, we were able to generate double transgenic mice to test whether *Kif11* overexpression prevents or reduces AD-related cognitive deficits in the 5xFAD mouse model. Additionally, we were able to test whether increased *Kif11* expression alone impacts learning and memory by comparing single transgenic Kif11OE mice to their wild-type (WT) littermates.

Kif11OE mice were mated with 5xFAD mice, and the progeny were backcrossed to obtain single and double transgenic mice in a >95% C57Bl/6J genetic background (see Methods). This generated litters consisting of non-transgenic WT littermates, single transgenic mice harboring the Kif11OE transgene (Kif11OE), single transgenic mice harboring the 5xFAD transgene (5xFAD), and double transgenic mice harboring both the 5xFAD and Kif11OE transgenes (5xFAD-Kif11OE). To determine whether 5xFAD-Kif11OE mice had altered expression of either the Kif11OE or the 5xFAD transgenes, we compared the mRNA expression levels of human *APP* and mouse *Kif11* in whole brain homogenates from 8-month-old mice. We found that both Kif11OE and 5xFAD-Kif11OE mice showed similar levels of *Kif11* overexpression (Figure 1A), and that the 5xFAD and 5xFAD-Kif11OE mice showed similar levels of *APP* overexpression (Figure 1B). These data demonstrate that mating Kif11OE mice successfully produced a viable double transgenic mouse model of AD that overexpress both mouse *Kif11* and human *APP* at levels similar to those detected in their respective single transgenic littermates.

**Figure 1:**
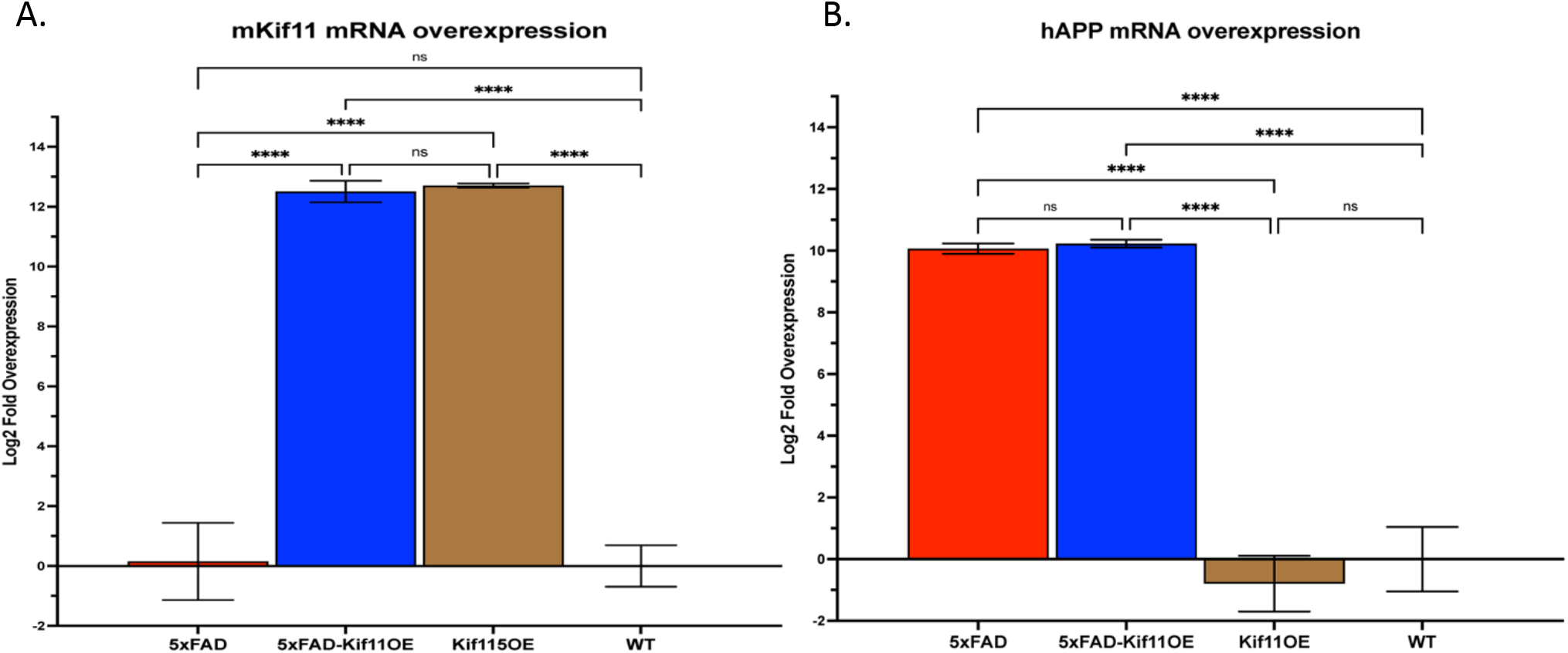
Quantitation of human *APP* and mouse *Kif11* mRNA expression levels in whole brain tissue from 5xFAD, 5xFAD-Kif11OE, and Kif11OE mice and their WT littermates. Quantitative PCR (qPCR) was used to determine mRNA expression levels of (A) human *APP* (*hAPP*) and (B) mouse *Kif11* (*mKif11*) relative to mouse *GAPDH*. Overexpression levels were calculated using the log2 fold change of the ΔΔCt. For quantification, the average expression level for each sample was calculated by averaging three technical replicates each. 5xFAD mice, N = 4 mice; 5xFAD-Kif11OE, N = 5 mice; WT, N = 3 mice; and Kif11OE, N = 7 mice.

### Increased *Kif11* expression rescues spatial and working memory deficits in 6-to 8-month-old 5xFAD mice

The onset of behavioral deficits in 5xFAD mice occurs at approximately five months of age, and the deficits increase with age (Richard et al., 2015, Jawhar et al., 2012, Oakley et al., 2006, Xiao et al., 2015), whereas motor deficits that could potentially confound behavioral testing do not occur until 9-12 months of age (Jawhar et al., 2012, O’Leary et al., 2020, O’Leary et al., 2018). Here, we focused on the performance of 6-to 8-month-old mice in the radial arm water maze (RAWM) task as a measure of spatial and working memory (Alamed et al., 2006, Boyd et al., 2010, Arendash et al., 2001). As the RAWM task is motor function-dependent, using this age group allowed us to measure robust effects of Aβ-induced cognitive deficits prior to the development of motor deficits in the 5xFAD mice (O’Leary et al., 2020, O’Leary et al., 2018).

The mice were first placed on the platform to learn its location within the room. The mice were then placed in one of the other five arms of the water maze with the goal of swimming to the location of a submerged platform, using only the environmental cues from the maze and the room (Figure 2A). The ability of the mice to learn the location of the submerged platform and escape the water was measured over two days in 30 trials of testing with 15 trials per day (Figure 2B). To control for the distance of the platform from the point of placement as well as for potential differences in the allocentric and egocentric abilities of the mice, the performance of each mouse was quantified in blocks of trials. The blocks consisted of six trials in the first two blocks of each day and three trials in the last block of each day (Figure 2B), with each trial beginning at a different placement arm (Figure 2B). Profiles of latency (time to reach the submerged platform) and errors (entry of the mouse into the incorrect arm) revealed that the 5xFAD-Kif11OE and Kif11OE groups performed more similarly to the WT group, and that the latency and error profiles of the 5xFAD group were markedly different from those of the WT, 5xFAD-Kif11OE, and Kif11OE groups (Figure 2C-D). Furthermore, quantification and comparison of latency profiles revealed that the 5xFAD mice showed a significantly longer latency compared to WT, Kif11OE, and 5xFAD-Kif11OE mice, and that the latency of 5xFAD-Kif11OE mice was not significantly different from that of WT or Kif11OE mice (Figure 2E). These findings indicate that increased expression of *Kif11* alone does not lead to motor deficits that affect RAWM latency compared to WT mice, and that increased *Kif11* expression leads to decreased RAWM latency in 5xFAD mice to levels similar to those of WT mice. There were no significant differences in the average number of errors per block made by any of the four groups during day 1 (Figure 2F). However, during day 2, the 5xFAD mice made significantly more errors compared to WT, Kif11OE, and 5xFAD-Kif11OE mice (Figure 2G). These findings show that increased expression of *Kif11* reduces the error rate of 5xFAD mice in the RAWM to levels similar to those of WT mice (Figure 2G). The fact that 5xFAD mice made significantly more errors on day 2 of testing, but not on day 1, indicates that 5xFAD mice have deficits in memory-dependent performance that are rescued by increased expression of *Kif11*. Taken together, our results show that increased expression of *Kif11* prevents deficits in learning and memory in the 5xFAD mouse model.

**Figure 2:**
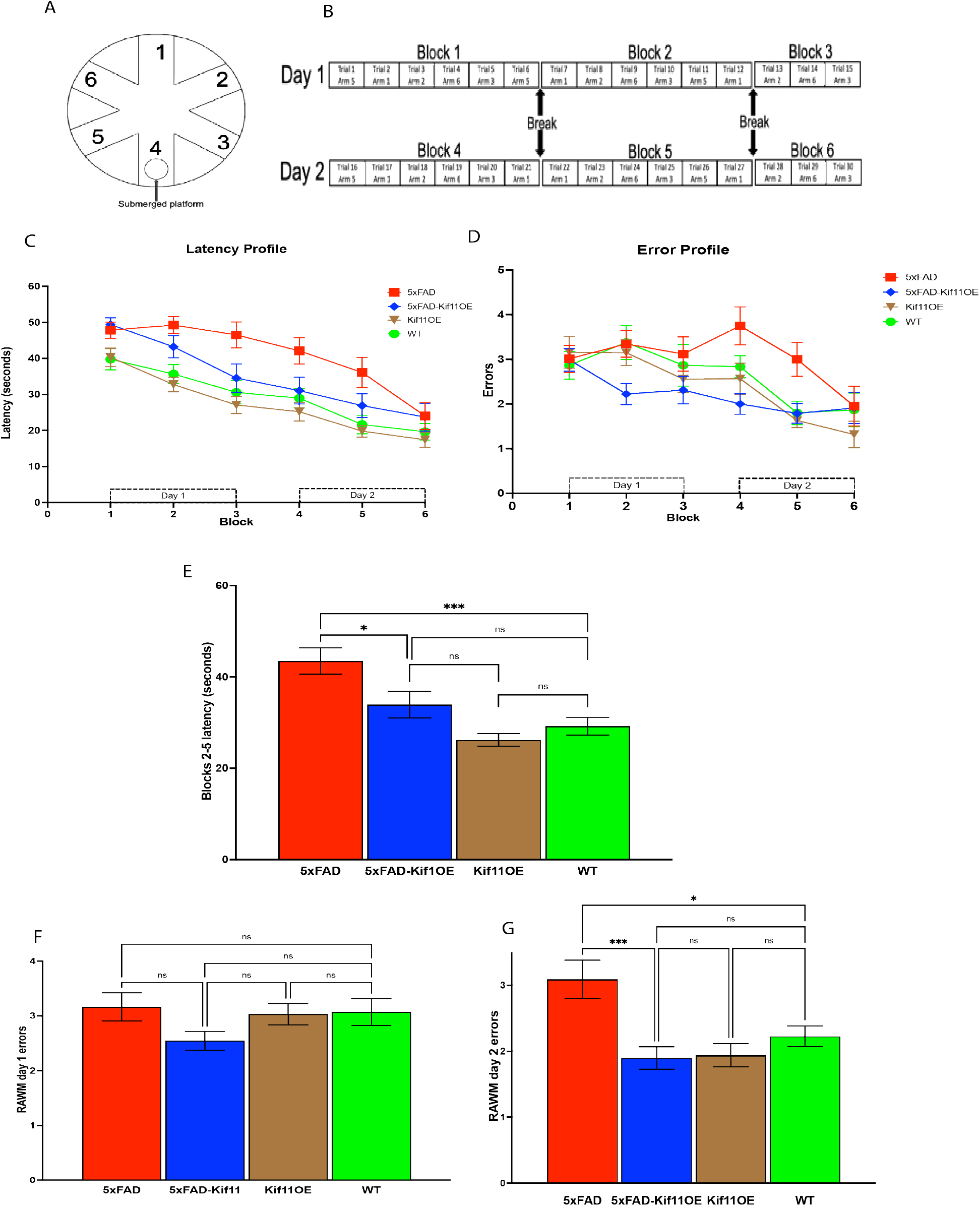
Increased *Kif11* expression rescues working and spatial memory deficits in 6-to 8-month-old 5xFAD mice in the radial arm water maze task (RAWM). (A) Schematic showing the six arms in the RAWM. (B) Illustration of arm placement and timing of breaks between the 30 trials in the RAWM during two days of testing with 15 trials/day. (C) Average time per block each group took to find the escape platform. (D) Average number of errors for each mouse at each block of testing. (E) Average latency performance in blocks 2–5. (F) Average errors made by each mouse on day one of testing. (G) Average errors made by each mouse on day two of testing. 5xFAD, N = 15; 5xFAD-Kif11OE, N= 15; Kif11OE, N = 19; and WT, N = 15.

### Increased *Kif11* expression rescues Aβ-mediated decreases in LTP

Hippocampal LTP has been postulated to be the cellular basis for learning and memory (Bliss and Collingridge, 1993, Roman et al., 1987). As cognitive decline is apparent in AD and in mouse models thereof, including 5xFAD mice, deficits in hippocampal LTP have been attributed to Aβ toxicity (Kimura and Ohno, 2009, Crouzin et al., 2013). We previously reported that Aβ-mediated inhibition of Kif11 is one potential mechanism by which Aβ inhibits hippocampal LTP (Freund et al., 2016). Because performance in the RAWM task depends on hippocampal function, we hypothesized that increased *Kif11* expression might help to maintain hippocampal LTP in 5xFAD mice in accordance with the prevention of the spatial learning and memory deficits we observed above. Specifically, we predicted that *Kif11* overexpression would prevent or reduce Aβ-mediated deficits in LTP.

As shown in Figure 3, extracellular recordings of synaptic responses measured as the initial slope of the field excitatory post-synaptic potential (fEPSP) at CA1 synapses of 6-to 8-month-old mice revealed that the fEPSP responses were markedly increased in the Kif11OE, WT, and 5xFAD-Kif11OE groups following the induction of LTP by delivery of two high-frequency stimulus (HFS) trains. In contrast to these results, the 5xFAD group fEPSP responses after HFS remained more similar to baseline responses recorded prior to HFS (representative traces are shown in Figure 3A) consistent with impaired LTP. Accordingly, at 60 min post-HFS, the 5xFAD group showed significantly less LTP of the fEPSP response compared to the WT group, whereas the 5xFAD-Kif11OE group showed no significant difference compared to either the WT or Kif11OE groups (Figure 3B-C). Our findings reveal that *Kif11* overexpression alone does not affect hippocampal LTP compared to age-matched WT mice, and, more importantly, that *Kif11* overexpression protects against Aβ-mediated deficits in LTP in 5xFAD mice, as shown in the 5xFAD-Kif11OE mice (Figure 3C). The observation that the Kif11OE mice performed similar to WT mice in the RAWM task and expressed similar levels of LTP suggests that the rescue of the cognitive deficits in 5xFAD-Kif11OE could be due to a compensatory interaction between the overexpressed Kif11 and Aβ overexpression that results from the FAD transgenes, as we predicted.

**Figure 3:**
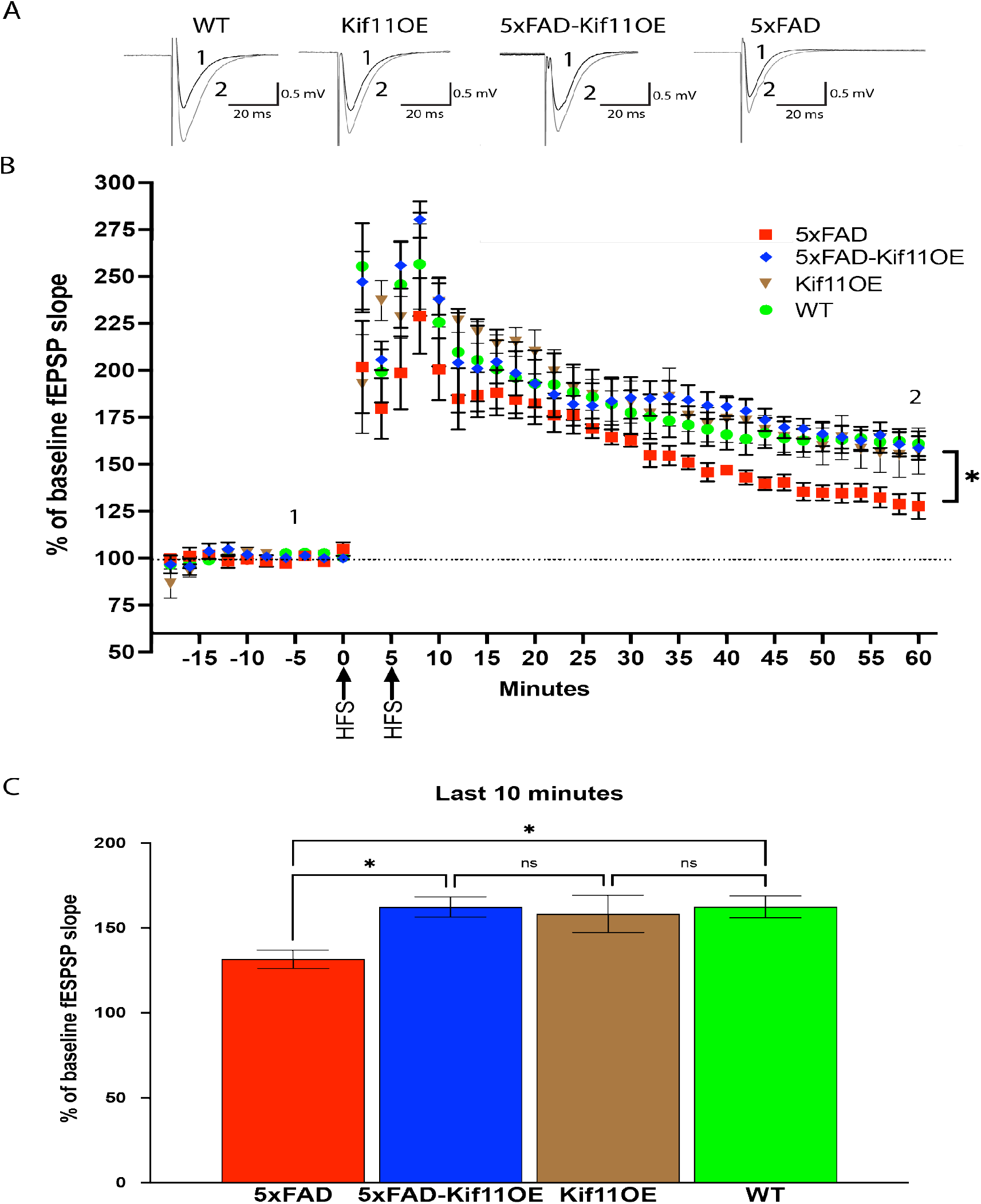
Increased Kif11 expression prevents decreases in early phase long-term potentiation (LTP) in 5xFAD mice. (A) Typical field excitatory postsynaptic potentials (fEPSPs) for each of the four groups of mice (5xFAD, 5xFAD-Kif11OE, Kif11OE, and WT) before (1, gray line) and 60 min after delivery of two 100 Hz, 1 second high frequency stimulus (HFS) trains delivered 5 min apart to induce LTP (2, black line). (B) Time course of fEPSP slope measurements (normalized as % of baseline) before and after two HFS trains (black arrows: 1 x 100 Hz each, 5 min apart). Data represent the mean ± S.E.M for five slices from four animals for the 5xFAD group, six slices from four animals for the 5xFAD-Kif11OE group, and five slices from three animals for the Kif11OE and WT groups (C) LTP normalized as % of baseline fEPSP slope at 50-60 min after HFS. Statistically significant differences between groups were determined by ordinary one-way ANOVA with post-hoc Šídák multiple comparison analysis. *P < 0.05.

These data suggest that Kif11 plays a key role in maintaining neuronal functions critical for learning and memory, and that the improved performance of the 5xFAD-Kif11OE mice compared to the 5xFAD mice in the RAWM can be attributed, at least in part, to the maintenance of hippocampal LTP mediated by increased expression of *Kif11*. These data provide further mechanistic support for the conclusion that Aβ-mediated inhibition of Kif11 contributes to cognitive dysfunction in AD.

### Increased expression of *Kif11* does not affect brain amyloid deposition in 5xFAD mice

The prevention or reversal of amyloid deposition by active or passive immune therapy against Aβ, by genetic removal of the gene encoding APOE (which is essential for Aβ polymerization), or by various immune modulators all lead to cognitive benefits in animal models and in at least some human trials compared to placebo, suggesting the targeting of amyloid as one approach to AD therapy. To test whether the rescue of AD phenotypes that we observed in our behavioral and electrophysiological experiments in 5xFAD-Kif11OE mice might similarly be the result of reduced amyloid deposition, we visualized Aβ accumulation in different brain regions in 5xFAD, 5xFAD-Kif11OE, and Kif11OE mice using both the NIAD4 amyloid-binding dye that recognizes beta-sheet structures (Nesterov et al., 2005) and the 6E10 antibody raised against amino acids 1-16 of Aβ. Histological staining of sagittal slices from 8-month-old mouse brains revealed that both 5xFAD mice (Figure 4A) and 5xFAD-Kif11OE mice (Figure 4B) had robust deposition of amyloid plaques, whereas the Kif11OE mouse had no amyloid signal (Figure 4C). Quantification of the fluorescence-positive area in multiple brain regions revealed that 5xFAD and 5xFAD-Kif11OE mice had similar levels of amyloid plaque burden, and Kif11OE mice lacked amyloid plaques, based on NIAD4 staining (Figure 4D) and 6E10 staining (Figure 4E). Furthermore, whole brain quantification of signal from both NIAD4 staining and 6E10 staining also revealed similar results, where the 5xFAD and the 5xFAD-Kif11OE mice showed similar levels of amyloid deposition, and the Kif11OE mice lacked amyloid plaques (Figure 5A-B). We did not observe any significant differences in DAPI staining in the same brain regions (Figure 6A) or with whole brain quantification of DAPI staining (Figure 6B) in 5xFAD, 5xFAD-Kif11OE, and Kif11OE mice. These data demonstrate that the rescue of learning and memory deficits and the maintenance of LTP in 5xFAD-Kif11OE mice is not due to reduced Aβ deposition. Instead, increased *Kif11* expression appears to bolster brain function and resiliency despite the presence of amyloid deposits in the brain.

**Figure 4:**
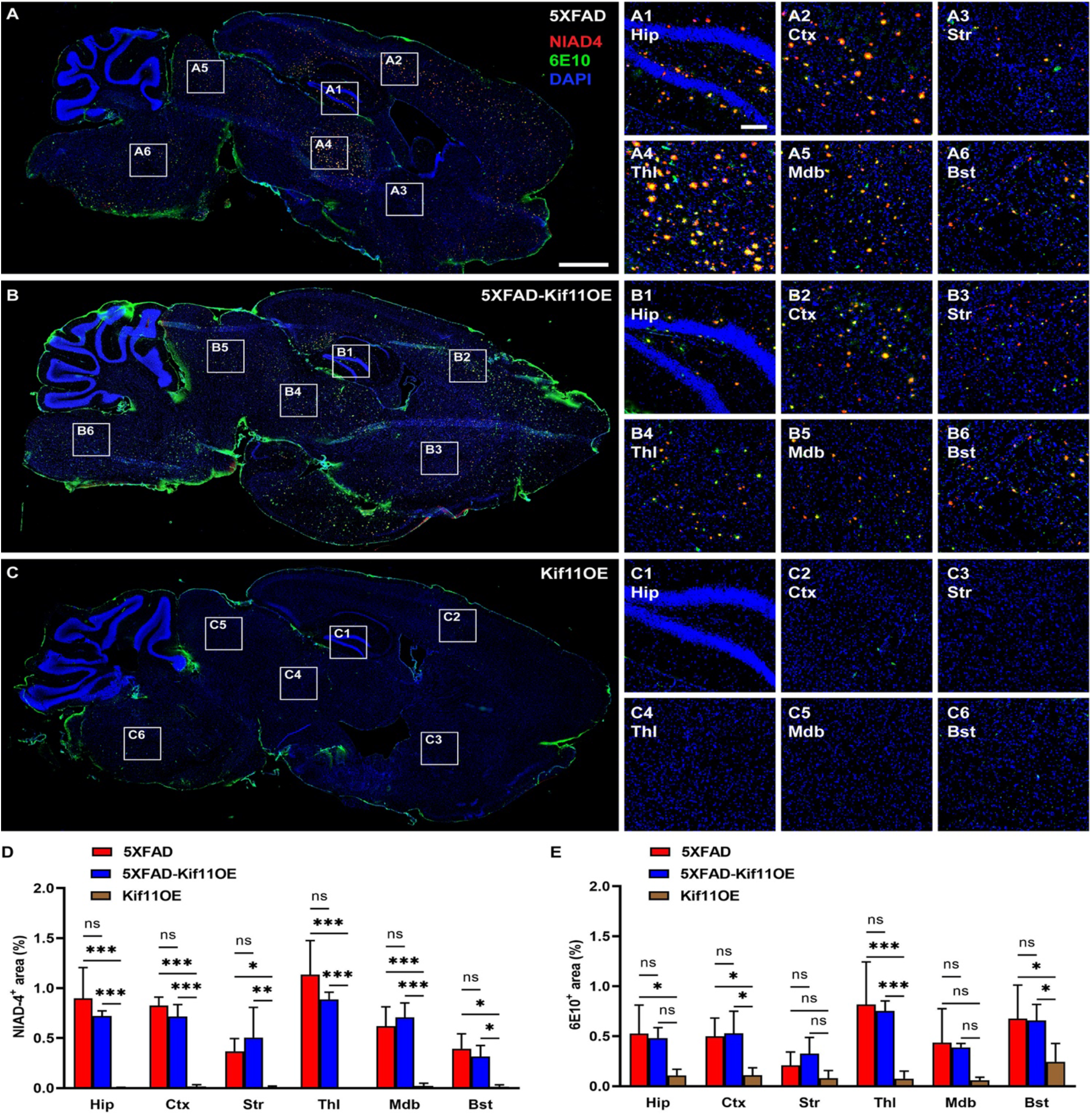
Increased expression of *Kif11* does not reduce amyloid plaques in the brains of 5xFAD mice. Shown are representative images of 20 μm-thick sagittal sections of mouse brain showing areas of the hippocampus (Hip), cortex (Ctx), striatum (Str), thalamus (Thl), midbrain (Mdb), and the brainstem (Bst) that were analyzed to detect amyloid using both NIAD4 staining (red) and the 6E10 antibody (green) and cell nuclei using DAPI (blue) in 5xFAD (A, A1-A6), 5xFAD-Kif11OE (B, B1-B6), and Kif11OE (C, C1-C6) mice. Quantification of the percent area positive for (D) NIAD4 staining or (E) 6E10 staining in the different brain regions in 5xFAD, 5xFAD-Kif11OE, and Kif11OE mice. Data represent the mean ± S.D. of N = 2–5 slices from three mice in each group. Statistical significance was calculated through ordinary one-way ANOVA with post-hoc Holm-Šídák’s multiple comparisons test. \**P* < 0.05, \*\**P* < 0.01, \*\*\**P* < 0.001.

**Figure 5:**
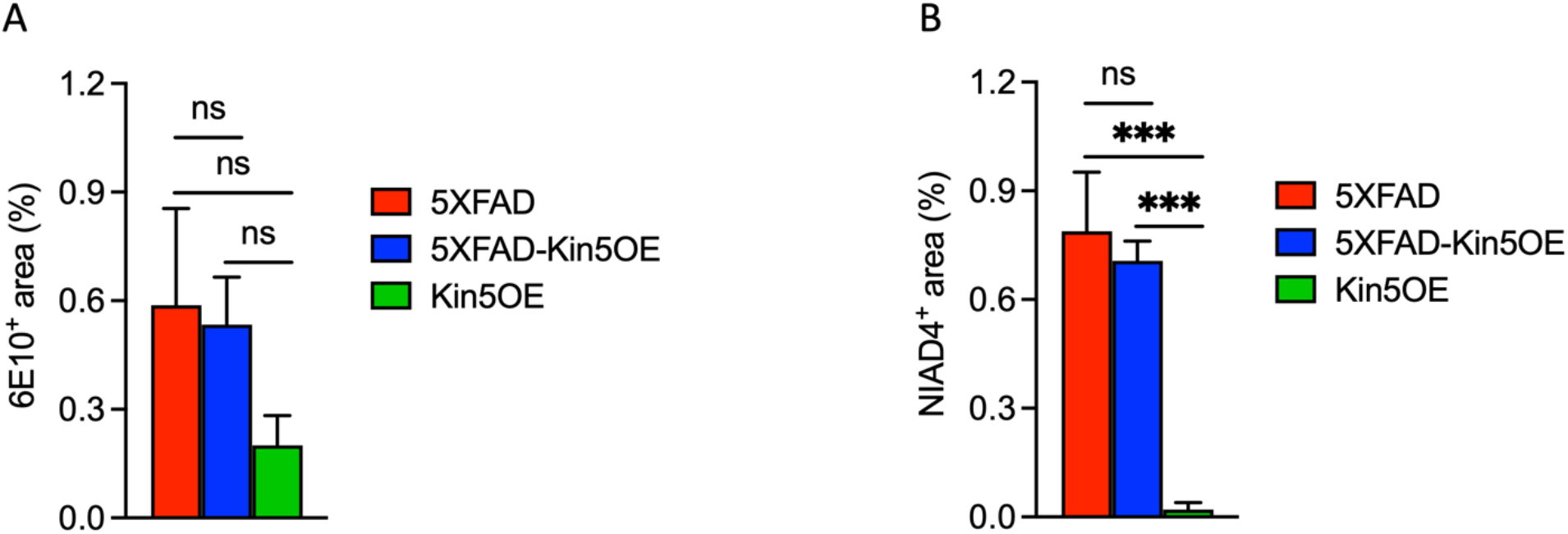
Amyloid burden in 6-to 8-month-old mice. Quantification of the percent area of 20 μm-thick sagittal sections of mouse brain slice from 5xFAD, 5xFAD-Kif11OE, and KIF11OE mice that were (A) positive for NIAD4 staining, (B) positive for 6E10 staining. Data represent the mean ± S.D. of N = 3 mice per group and 2–5 brain slices from each mouse. Statistical significance was calculated through ordinary one-way ANOVA with post-hoc Holm-Šídák’s multiple comparisons test. \*\*\**P* < 0.0001.

**Figure 6:**
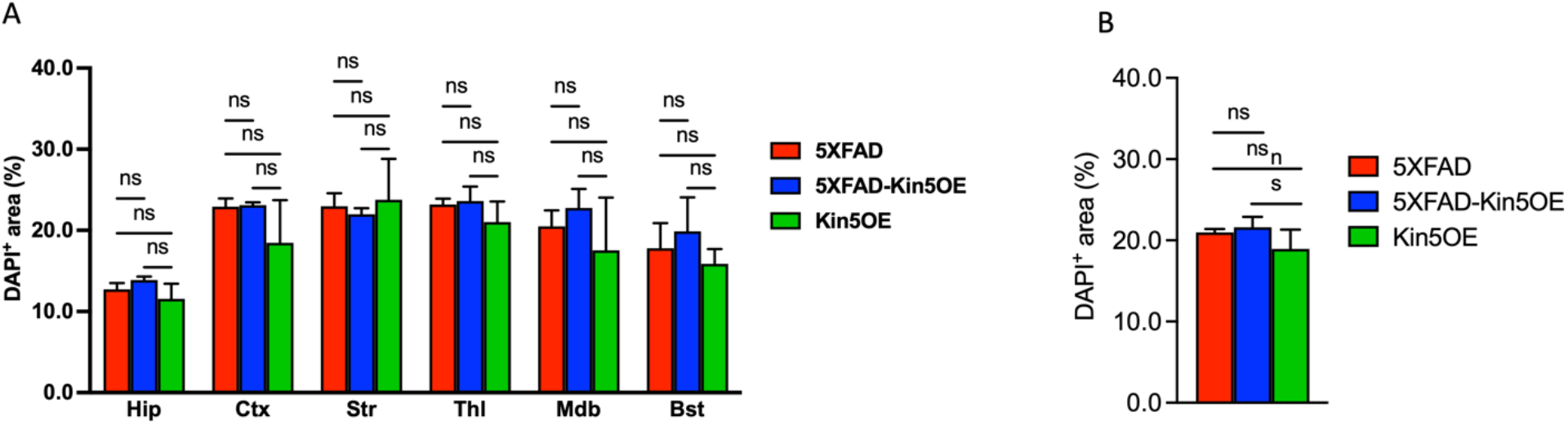
Percent area positive for DAPI staining in brains of 6-to 8-month-old mice. Quantification of percent area positive for DAPI staining in 20 μm-thick sagittal sections of mouse brain slices from 5xFAD, 5xFAD-Kif11OE, and KIF11OE mice. (A) Percent area positive for DAPI staining in specific brain regions including the hippocampus (Hip), cortex (Ctx), striatum (Str), thalamus (Thl), midbrain (Mdb), and brainstem (Bst). (B) Percent area positive for DAPI staining in whole 20 μm-thick sagittal sections of mouse brain slices. Data represent the mean ± S.D. of N = 3 mice per group and 2–5 brain slices from each mouse. Statistical significance was calculated through ordinary one-way ANOVA with post-hoc Holm-Šídák’s multiple comparisons test. \*\*\**P* < 0.0001.

### Kif11 prevents Aβ-mediated loss of spine density in primary rat neurons

Aβ toxicity has been attributed to synaptic loss in AD patients as well as in cell and mouse models of AD (Ingelsson et al., 2004, Spires et al., 2005, Freund et al., 2016, Hsieh et al., 2006), DeKosky and Scheff, 1990, Scheff et al., 2007). Considering the role of Kif11 in maintaining neuronal morphology and structure, and in view of our previous finding that Aβ-mediated inhibition of KIF11 is one mechanism likely to contribute to dendritic spine loss in AD (Freund et al., 2016), we sought to determine whether overexpression of Kif11 can block Aβ-mediated spine loss. To this end, we employed a cell culture model that overexpresses Kif11 using transient transfection to determine whether acute Kif11 overexpression alone impacts spine density. Transient transfection and overexpression of Kif11 in rat primary neurons led to a dosedependent decrease in spine density with increasing concentrations of added Kif11-expressing plasmid (Figure 7). When combined with our previous findings showing that Kif11 inhibition with monastrol also reduces dendritic spine density, these findings for Kif11 overexpression indicate that Kif11 activity likely has to be maintained in a narrow range to support its normal cellular functions. These findings also clearly illustrate the structural role of Kif11 in dendrites and in dendritic spines and are in agreement with previous studies showing that Kif11 regulates neuronal architecture (Freixo et al., 2018, Kahn et al., 2015, Yoon et al., 2005). In parallel, we also employed an AD-Kif11OE cell culture model system in which primary rat neurons were transiently transfected to express human *APP* harboring the Swedish mutation (APPswe) (Hsieh et al., 2006), which results in Aβ overproduction and FAD/EOAD in humans (Mullan et al., 1992), with or without Kif11 overexpression. We hypothesized that the effects of Aβ on dendritic spines, which are important for learning and memory (Spires et al., 2005, Wei et al., 2010), would be reduced by acute *Kif11* overexpression. In agreement with previously published reports, we found that transfection of APPswe alone led to significant spine loss, which can be attributed to Aβ production and toxicity. In contrast, co-transfection of the APPswe vector together with a low amount of the Kif11 overexpression vector, which produced only a small amount of spine loss compared to control on its own (Figure 7), was able to prevent dendritic spine loss from reaching the higher levels normally seen for APPswe transfection alone (Figure 8A-B). These data showing reduced APPswe-mediated spine loss in the presence of Kif11 overexpression support our hypothesis that Aβ-mediated spine loss is due in part to inhibition of Kif11 (Freund et al., 2016) and suggest a potential mechanism by which Kif11 overexpression helps to maintain normal cognitive function and hippocampal LTP in 5xFAD mice despite having no effect on Aβ production and amyloid deposition. Thus, based on our findings, we conclude that overexpression of Kif11 compensates for the inhibitory effects of Aβ on molecules important for supporting neuronal structures and functions required for learning and memory.

**Figure 7:**
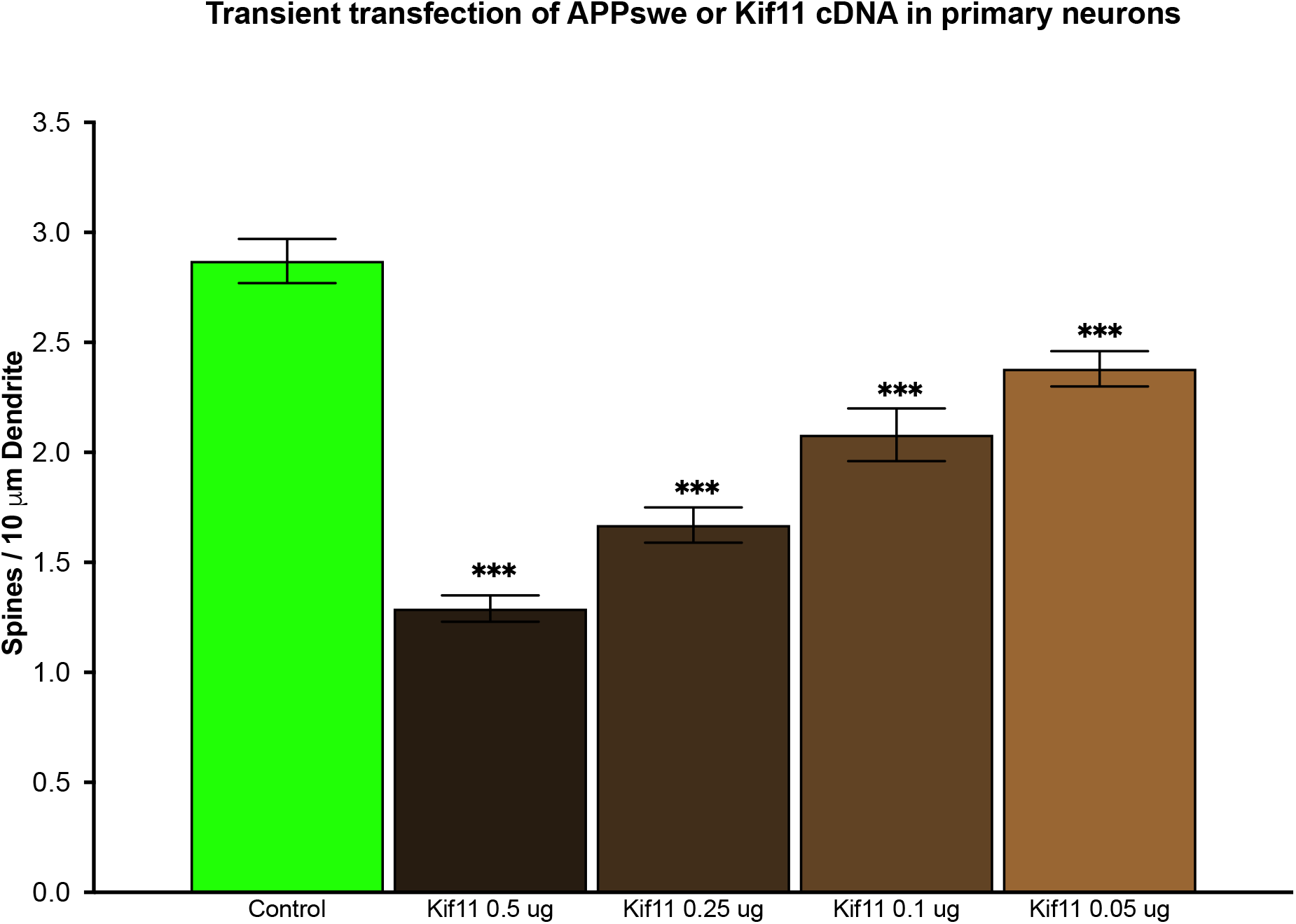
Spine density measurements in primary rat neurons transfected with different concentrations of the Kif11 overexpression plasmid. Spine density measurements from primary rat neurons transfected with a GFP plasmid alone (Control) or with a GFP plasmid and decreasing amounts of a Kif11 overexpression plasmid (0.5–0.05 μg). Statistical significance was calculated through ordinary one-way ANOVA with post-hoc Bonferroni’s multiple comparisons test. Asterisks above each bar indicate statistical significance compared to control. \*\*\**P* < 0.001.

**Figure 8:**
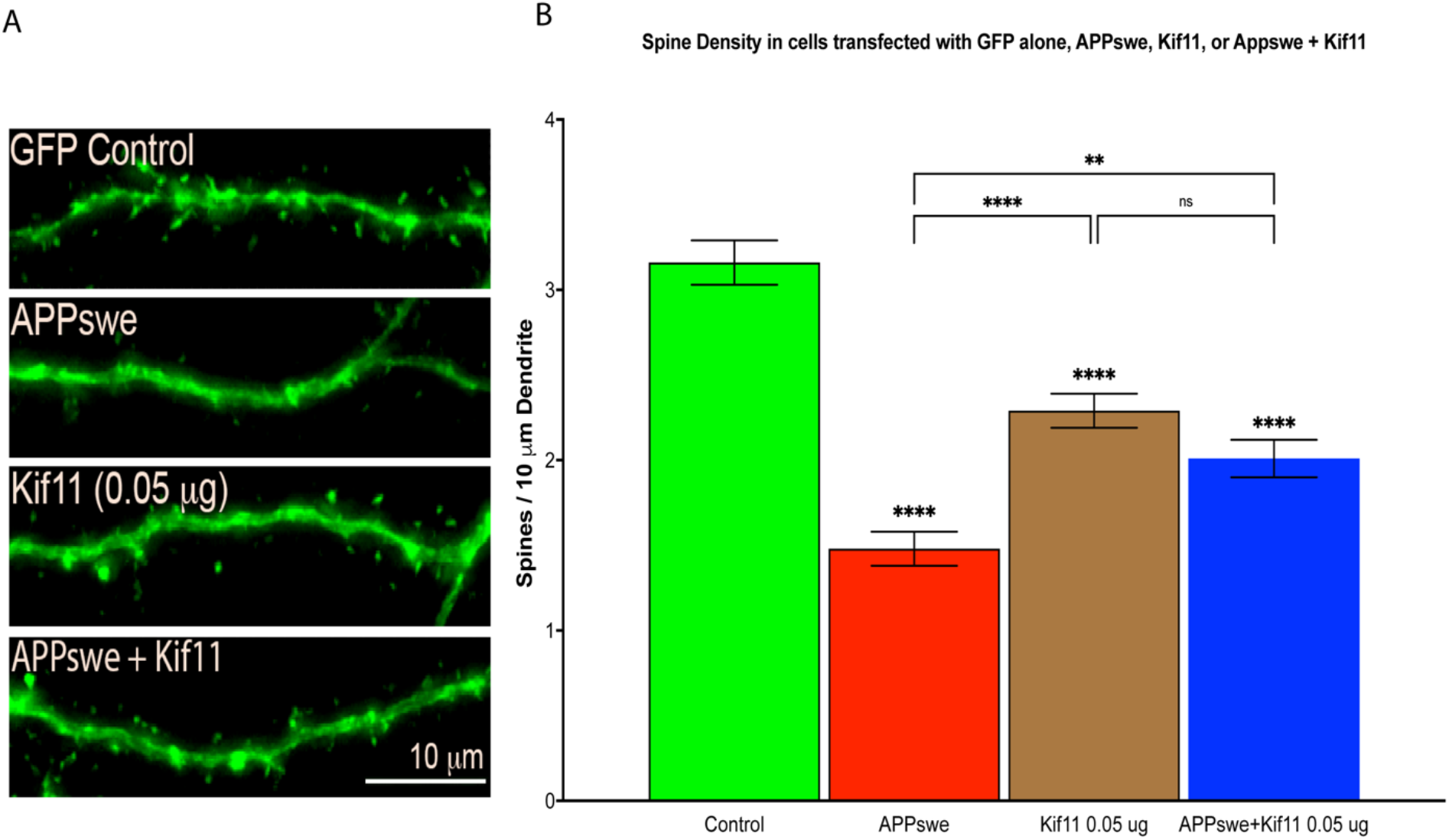
Moderate overexpression of Kif11 results in some dendritic spine loss but prevents more substantial mediated by Aβ-in primary rat neurons. (A) Representative images of primary rat neurons transfected with a plasmid to express GFP alone (Control, N = 28), with a plasmid to express GFP together with a plasmid to express the human APP gene harboring the “Swedish” double mutation (APPswe, N = 30), with 0.05 μg of a plasmid to express Kif11 (Kif11 0.05 μg, N = 30), or with 0.05 μg of a plasmid to express Kif11 together with the APPswe plasmid (Kif11 0.05 μg + APPswe, N = 28). (B) Spine density measurements from primary rat neurons transfected with GFP alone (Control), with GFP and APPswe (APPswe), with GFP and Kif11 (Kif11 0.05 μg), or with GFP, APPswe, and Kif11 (APPswe + Kif11 0.05 μg). Statistical significance measured by ordinary one-way ANOVA with post-hoc Šídák’s multiple comparisons test. Asterisks above each bar indicate statistical significance compared to control. **P<0.01, **** P < 0.0001

## Discussion

Many current experimental treatments for AD have focused on reducing Aβ production or on increasing the clearance of Aβ plaques (reviewed in (Imbimbo and Watling, 2019, Demattos et al., 2012)). Most of these approaches have failed to prevent or reverse cognitive decline in clinical trials, although one such anti-Aβ human monoclonal antibody drug (aducanumab/Aduhelm) may modestly slow cognitive decline compared to placebo (Graham et al., 2017, Cummings et al., 2014, Cummings et al., 2020). Clearly, additional approaches to the development of AD therapeutics are needed. Our work presents *in vitro* and *in vivo* experiments that together point towards a key role for the KIF11 microtubule motor in the maintenance of cognitive ability, wherein its inhibition by Aβ over the course of AD leads to cognitive deficits. Our behavioral and electrophysiological experiments show that increased *Kif11* expression overrides the inhibitory effects of Aβ, thereby preserving LTP and improving cognitive performance. The mechanism by which increased *Kif11* expression overrides the damaging effects of Aβ likely derives from protecting this important microtubule motor protein from complete inhibition by Aβ to preserve its many cytoskeletal functions that regulate neuronal morphology, including maintaining spine density, which has been previously hypothesized to serve as a marker of cognitive resilience in AD (Walker and Herskowitz, 2020, Boros et al., 2017). Our finding that *Kif11* overexpression does not affect Aβ production or deposition in the 5xFAD mice shows that the beneficial effects of *Kif11* overexpression on cognition and LTP in our combined *Kif11*OE-5xFAD mice occurred despite the persistence of amyloid plaques and pathology in the brains of 5xFAD-Kif11OE mice.

Our data and experimental models provide evidence for the importance of KIF11 in maintaining cognitive function and synaptic plasticity in the presence of Aβ and elucidate the potential for novel AD therapeutic strategies that target Aβ toxicity based on enhancing KIF11 activity. In particular, we propose that overriding the toxic effects of Aβ by either increasing *Kif11* mRNA expression levels or blocking Aβ-mediated inhibition of KIF11 may be a viable therapeutic strategy for the treatment of AD.

As a whole, the data we present herein further highlight the importance of KIF11 in learning and memory and the possibility of developing therapies for AD that focus on maintaining neuronal structures and functions by targeting molecules involved in microtubule dynamics and stability. Although chemical compounds that alter microtubule dynamics were investigated previously as a potential therapy for AD (Zhang et al., 2012, Brunden et al., 2010, Brunden et al., 2014), our data presented here demonstrate the potential efficacy of targeting a microtubule motor protein inhibited by Aβ, specifically KIF11, and its increased expression and/or activity as a potential therapeutic for AD. To our knowledge, there have been no other *in vivo* studies showing that increased expression of a gene such as *KIF11*, whose protein product is inhibited by Aβ and is involved in regulating microtubule dynamics and organization, offsets the cognitive deficits caused by Aβ. Thus, our findings illustrate the potential efficacy of a new approach to AD therapy that involves overriding or blocking the inhibitory effects of Aβ on a specific cellular protein.

It is of interest to note that, in addition to Aβ, excess Tau has also been shown to inhibit *Kif11* and consequently cause the cell cycle defects, chromosome mis-segregation, neuronal aneuploidy, and apoptosis that characterize a *Drosophila* model of frontotemporal dementia (FTD, also termed frontotemporal lobar degeneration, FTLD) (Bougé and Parmentier, 2016, Malmanche et al., 2017). Such microtubule-dependent cell cycle defects, including neuronal aneuploidy and apoptosis, have also been observed in human FTD/FTLD caused by mutations in the Tau/MAPT gene (Caneus et al., 2018) (for discussion, see (Potter et al., 2019)). Thus, two of the major pathological and biochemical features of AD — Aβ and Tau — both inhibit KIF11 activity and function, further supporting our conclusion that the microtubule motor protein KIF11 is a component in the AD pathogenic pathway and a potential focus of future therapeutic drug development efforts, with potential applicability to other neurodegenerative diseases.

### Transient transfection of APPswe or Kif11 cDNA in primary neurons

## Methods

Requests for reagent and resource sharing information and requests for resources and reagents should be directed to and fulfilled by the lead contact, Huntington Potter

### Animals

Male and female 5xFAD (Oakley et al., 2006) or *Kif11OE* (Tg*[Pim1-Eg5]Jus* mice) (Castillo et al., 2007), referred to herein as Kif11OE mice, were used for breeding and for generation of the 5xFAD-Kif11OE transgenic mouse strain. Both male and female 6-to 8-month-old mice were used for electrophysiology experiments and for behavioral tests in the RAWM. All animal experiments and husbandry were approved by and conducted in accordance with the Institutional Animal Care and Use Committee (IACUC) guidelines and with approval from the University of Colorado Anschutz Medical Campus.

### Experimental models

Fully congenic male and female C57Bl/6J mice hemizygous for the 5xFAD transgenes that contain three mutations in human *APP* (Swedish: K670N, M671L, Florida: I716V, and London: V717I) and two mutations in human *PSEN1* (M146L and L286V) (Oakley et al., 2006) were obtained from Jackson Labs (JAX MMRRC Stock No: 34848-JAX). These mice were used to establish an in-house breeding colony in which 5xFAD mice were bred to fully congenic WT C57Bl/6J mice obtained from Jackson labs (Jax Stock Number: 000664). Establishment of the 5xFAD breeding colony took place in-house and utilized both male and female mice bred with respective WT C57Bl/6J mice. To establish the Kif11OE colony, we obtained cryo-preserved sperm from fully congenic FVB/NJ male mice harboring the p*Pim1Eμ-Eg5* transgene (Kif115OE) from the laboratory of Dr. Monica Justice at The Hospital for Sick Children, Peter Gilganís Centre for Research and Learning, Toronto, Ontario, Canada. This mouse model utilizes the p*Pim1Eμ-Eg5* transgene where the lymphoid specific Eμ enhancer in tandem with the *Pim1* promoter drive the expression of mouse Kif11. The transgene also contains the MuLV long terminal repeat to further amplify expression. The *Kif11* transgene in this mouse model has been shown to result in an up to 60-fold increase in Kif11 expression levels in the brain (Castillo et al., 2007). After we received the cryo-preserved sperm, rederivation of the transgenic mouse line was carried out through *in vitro* fertilization (IVF) by the University of Colorado Anschutz Medical Campus Gates Bioengineering Core. Briefly, mouse oocytes were generated using a fully congenic WT C57Bl/6J female mouse, creating mixed background (C57Bl/6J-FVB/NJ) Kif11OE progeny. Kif11OE mice produced through the rederivation processes were bred strictly with non-related fully congenic C57Bl/6J WT mice to backcross the mice to a predominant C57Bl/6J background. Breeding took place in-house and utilized both male and female mice from each group. We continued to backcross the Kif11OE mice with fully congenic C57Bl/6J mice throughout the entirety of this study. To develop a 5xFAD Alzheimer’s mouse model that overexpresses mouse Kif11 (5xFAD-Kif11OE), we used unrelated male and female mice that carried either the 5xFAD transgenes or the Kif11OE transgene for breeding. Kif11OE mice from the F4-F9 C57Bl/6J backcross generation were then bred with fully congenic C57Bl/6J 5xFAD mice to create a mouse line that was predominantly on a C57Bl/Bl6J background (93%-99% C57Bl6/J). Only mice derived from the 5xFAD x Kif11OE cross were used in experiments.

### Genotyping of transgenic mouse lines

To obtain tissue samples for genotyping, small ear clips were taken from 21-to 28-week-old mice anesthetized with 4% isoflurane. At that time, the mice were also given a subcutaneous radio frequency radio identification (RFID) tag (Trovan, Ltd, cat: 040104) and weaned from the mother. DNA was extracted from the ear tissue using Extracta DNA Prep (Quantabio) according to manufacturer guidelines. Briefly, the tissue was incubated for 30 min at 95°C in 300 μl extraction reagent, cooled to room temperature, and 300 μl stabilization buffer was then added to the sample. PCR Genotyping of Kif11OE mice for the p*Pim1Eμ-Eg5* transgene was carried out using the protocol established by Castillo et. al., (2007). Primers specific to Kif11 (5’-TGACTTCCGATGAAGAAAGC-3’) and the MuLV LTR of the transgene construct (5’-GATACACGGGTACCCGGGCG-3’) were used with an annealing temperature of 58°C and 20 cycles of PCR. The resulting PCR products were then ran on a 1% agarose gel that revealed binary results, where samples that yielded a single ~1,000 bp band were considered to be positive for the *pPim1Eμ-Eg5* transgene, and samples that yielded no band were considered non-transgenic (WT). 5xFAD mice were genotyped according to guidelines established by Jackson Labs (https://www.jax.org/Protocol?stockNumber=008730&protocolID=34650). Briefly, a primer probe specific to the APPswe mutation (TmoIMR0076-Fluorophore-1 5’-CATTGGACTCATGGTGGGGGGGGGTG-3’) was paired with primers specific to the APPswe transgene in the 5xFAD mouse (5’-TGGGTTCAAACAAAGGTGCAA-3’, 5’-GATGACGATCACTGTCGCTATGAC-3’) and subjected to 40 cycles of RT-PCR with an annealing temperature of 60°C. Samples that showed amplification of the PCR product through real-time fluorescence after 20 cycles were considered positive for the 5xFAD transgene, and samples that did not show any amplification during the 40 cycles were considered to not contain the 5xFAD transgene. PCR genotyping for each mouse strain used PerfeCTa^®^ MultiPlex qPCR SuperMix, Low ROX™ (Quantabio, cat: 95063) at a 1X concentration.

### Gene expression analyses

Both *APP* and *Kif11* expression were analyzed using qRT-PCR. Whole brains were extracted from mice anesthetized with 100 μl pentobarbital. Once anesthetized, the mice were sacrificed using cervical dislocation and decapitation, and the brain was quickly removed and quartered. The quartered pieces of each brain were placed into individual RNAase-free 2 ml microfuge tubes and flash frozen with liquid nitrogen. Brain tissue samples were homogenized using ultrasonication in 1 ml of 35°C Trizol (Invitrogen cat: 15596018). The samples were incubated for 5 min at room temperature and vortexed for 10 sec intermittently during the incubation. To extract the RNA, 200 μl chloroform was added to each sample, followed by vortexing for 15 sec, incubation for 1 min at room temperature, vortexing for 15 sec, and centrifugation at 13,000 x g for 10 min. The supernatant was then collected for the isolation and purification of the RNA using the RNeasy Mini Kit (QIAGEN, cat: 74104) according to manufacturer instructions. After the RNA was eluted, cDNA was synthesized using the iScript™ cDNA synthesis Kit (Bio-Rad). Expression levels of *Kif11* and *APP* were then measured by multiplex qRT-PCR using mouse *Kif11* and human *APP* primer probes (Applied Biosystems, Mm01204225_m1, Cat#4448489 and Hs0016908_m1 Cat#4331182, respectively) in combination with mouse *GAPDH* primer probes (Applied Biosystems Cat#4331182/AssayID-Mm99999915_g1) for normalization. Expression levels were calculated as the log2-fold change of the ΔΔCt.

### Radial Arm Water Maze (RAWM)

We measured working memory in our mouse models using the performance of each mouse in a six-arm RAWM. Testing was carried out using methods described previously by Alamed et. al., (2006) with minor modifications. We used modular plastic inserts placed inside a water tank to create six radially distributed swim arms emanating from a central circular swim area. The tank was placed on a table so that the top of the tank was approximately 6.5 feet from the ground. This allowed the test to take place without visual cues from the surrounding room environment and eliminated influence of the performance of the mouse that might be caused by activities of the people administrating the test. Visual cues to facilitate spatial awareness of the mouse were not added to the maze. The tank was filled with water the night prior to testing to allow the water to reach room temperature, and the water was dyed white with non-toxic watersoluble paint on the morning of testing. An escape platform made of translucent plexiglass was placed in one of the six arms and remained in that same arm throughout the testing. Prior to the beginning of testing on day one of the two-day-long test, the mouse was introduced to the maze by placing it on the submerged platform and allowing it to remain there for 60 sec. Afterwards, the mouse was placed in its holding cage until the beginning of the test. Mice were tested in groups of 2-4 mice per block of testing, with each mouse per group undergoing the test sequentially, which allowed each mouse to rest between trials during that block of testing. After the mice from each group had completed each block of trials, each mouse was then placed in one of thk/bfhe five arms that did not contain the escape platform and was given 60 sec to complete the maze. If the mouse failed to complete the maze in 60 sec, the mouse was guided to the platform with a wooden guiding board and allowed to place itself on the platform where it remained for 60 sec. Each mouse underwent two days of testing with 15 trials per day. The performance of each mouse was measured using both the time (latency) and the number of errors the mouse made during testing. Errors were considered when the mouse fully entered (i.e., entire body) an arm that did not contain the escape platform or when the mouse entered the correct arm with the escape platform but did not swim to or locate the escape platform. Measurements of performance relative to the average performance of the WT group were calculated as the percent difference of each mouse at each block compared to the average performance of the WT group at each block. Statistical analyses for RAWM performance used GraphPad Prism software version 9.0.2 (https://www.graphpad.com/scientific-software/prism/).

### Electrophysiology

Electrophysiology recordings for LTP measurements were carried out on transverse 400 μm-thick hippocampal slices as described previously (Freund et al., 2016) from mice that had not undergone behavioral testing in the RAWM. All of the mice used for electrophysiology experiments were progeny of 5xFAD x Kif11OE breeding. Mice were anesthetized with isoflurane, decapitated, and the brain was removed. To cool the interior of the brain, the entire brain was placed in ice-cold cutting solution containing 220 mM sucrose, 25 mM D-Glucose, 2 mM KCl, 12 mM MgCl_2_, 0.2 mM CaCl_2_, 1.25 mM NaH_2_PO_4_, and 26 mM NaHCO_3_ for 60 sec. Afterwards, the hippocampi were dissected from the brain and sliced transversely with a McIlwain tissue slicer. Once the slices were made, they were placed in a recovery chamber containing artificial cerebral spinal fluid (aCSF) (124 mM NaCl, 11 mM D-glucose, 3.5 mM KCl, 1.3 mM MgCl_2_, 2.5 mM CaCl_2_, and 25.9 mM NaHCO_3_) for at least 1 h. Individual hippocampal slices were then transferred to a recording chamber superfused at a bulk rate of 2-3 ml/min with 30°C aCSF. To measure fEPSP responses, we placed stimulus and recording electrodes in the collateral axon pathway of the CA1 dendritic field. Measurements of the fEPSP response were an average of three responses to stimuli delivered 20 sec apart. Prior to baseline recordings, we generated an input-output curve by increasing the stimulus voltage and recording the synaptic response until either a maximum response was reached, or until a population spike was observed in the fEPSP response. Stimulus intensity was set at 40–50% of the maximum stimulus intensity. Once proper stimulus intensity was determined, we took baseline recordings of the fEPSP response at a stimulus intensity specific to each slice. Baseline recordings were measured for 20 min, after which we administered HFS. HFS consisted of two trains of 100 Hz stimulation for 1 sec, with an intertrain interval of 5 min. Measurements of the fEPSP response to stimulus were monitored for 60 min after the first HFS was delivered. Statistical analyses for electrophysiology recordings used GraphPad Prism version 9.0.2 (https://www.graphpad.com/scientific-software/prism/).

### Neuronal cultures

Hippocampal neuronal cultures were prepared from postnatal day 0-2 male and female Sprague-Dawley rats, plated at medium density (300-450 cells/mm^2^) on glass coverslips and maintained in Neurobasal plus B27 (Invitrogen) and GlutaMAX (Invitrogen) medium until transfection on day-*in-vitro* (DIV) 11-12 using Lipofectamine 2000 (Invitrogen) as described previously (Keith et al., 2012, Freund et al., 2016) with plasmids encoding green fluorescent protein (GFP) (pEGFPN1; Clontech), mouse *Kif11* (Myers and Baas, 2007), or the human APPswe mutant (Hsieh et al., 2006). On DIV 14-15 (three days post-transfection), neurons were fixed in 4% paraformaldehyde, and the coverslips were mounted on slides with Pro-Long Gold (Invitrogen). Images of dendrites in GFP-transfected neurons were acquired on an Axiovert 200M microscope (Zeiss) with a 63X objective (1.4NA, plan-Apo), a 1.5X magnifier, and a CoolSNAP2 (Photometrics) CCD camera. Focal plane Z-stacks (0.5 micron sections) were acquired, deconvolved to correct for out-of-focus light, and 2D maximum intensity projections generated (Slidebook 5.0-6.0, Intelligent Imaging Innovations). Spine numbers were quantified from projection images using the ruler tool in Slidebook 5.0-6.0 software with manual counting and were expressed as the number of spines/10 μm of dendrite using measurements obtained for multiple lengths of dendrites (N = the number of lengths of dendrite) taken from multiple images across three independent neuronal cultures for each experimental treatment condition. Statistical analyses for dendritic spine density used GraphPad Prism version 9.0.2 (https://www.graphpad.com/scientific-software/prism/).

### Mouse brain collection and amyloid staining

To determine amyloid plaque burden in our mouse models, we used 20 μm-thick sagittal cryo-sectioned mouse brains. Prior to cryosectioning, the mice were anesthetized with 100 μl of pentobarbital. Once fully anesthetized, transcardial perfusion was performed with 0.9% NaCl in H_2_O. A Leica Biosystems Perfusion Two™ automated pressure prefusion system (cat: 39471005) was used to circulate the saline solution. Mice were perfused until the liver of the mouse showed evidence of being clear of blood or for 5 min after the start of perfusion. After perfusion, the brain was removed rapidly, and the hemispheres were separated. The two hemispheres were then placed in 10 ml of 4% paraformaldehyde in PBS for 72 h. The brains were then transferred into a 20% sucrose solution for 24 h and placed in a 30% sucrose solution and stored at 4°C until slicing. At 12-24 h before slicing, the brains were removed from the sucrose solution and placed in a 50 ml conical tube and stored at −80°C. On the day of cryosectioning, one hemisphere of the brain was placed inside the cryostat for 1 h to allow for acclimation to the −20°C environment. The brain hemisphere was then embedded in chilled Tissue Tek^®^ O.C.T™ compound, and incubated for 30 min inside the cryostat. Sagittal sections of the brain were obtained by slicing the brain at 20 μm laterally from the midline. Each slice was then placed onto a Fisherbrand^®^ Superfrost™ Plus Microscope Slide (cat: 12-550-15) and placed into a slide box on dry ice and stored at −80°C until staining.

For amyloid staining, tissue sections were permeabilized with 0.1% Triton X-100 in DPBS for 15 min and then blocked with 3% bovine serum albumin (BSA) in DPBS for 90 min at room temperature. Tissue sections were incubated with mouse anti-Aβ primary antibody (6E10, BioLegend #803014, 1:250) in 3% BSA in DPBS overnight at 4°C, followed by incubation with Alexa Fluor^®^ Plus 488-conjugated goat anti-mouse IgG secondary antibody (Invitrogen #A32723, 1:500) in 3% BSA in DPBS for 45 min at room temperature. Tissue sections were then stained with the amyloid-binding dye NIAD-4 (Cayman Chemical #18520) at 10 μM in DPBS for 10 min at room temperature, and the nuclei were stained using Hoechst 33342 (Thermo Scientific #62249) at 1 μg/ml in DPBS for 10 min at room temperature. Slides were imaged on an Olympus IX83 inverted fluorescence microscope at 10X magnification and then analyzed using Cell Sens v1.12 software (Olympus). Data from 2-5 tissue sections were averaged for each mouse.

## Acknowledgements

We would like to thank Dr. Jennifer Sanderson for help in preparing for the electrophysiology experiments. Additionally, we would like to thank Neil Markham for his mentorship, contributions, and help in the maintenance of our mouse colony. We would also like to thank Vanesa Adame for her help and contributions to the behavioral studies presented here. We thank the CU Anschutz Neurotechnology Center Animal Behavior Core staff, specifically Drs. Michael Mesches and Nicolas Busquet for their help and contributions to the mouse behavior studies. Finally, we would thank entire staff of the CU Anschutz Office of Live Animal Research for their care and maintenance of our mouse colonies. This work was supported by NIH grants F99NS115330 to E.M.L and R01NS110383 to M.L.D. as well as gift funds to the University of Colorado Alzheimer’s and Cognition Center and H.P.

